# Evolutionary-scale protein language models uncover beneficial variants in a Sorghum bicolor diversity panel

**DOI:** 10.64898/2026.04.10.717708

**Authors:** Natasha H. Johansen, Janek Sven-Ole Sendowski, Eleni Nikolaidou, Savvas Chatzivasileiou, Shuai Wang, Baoxing Song, Andrew Olson, Thomas Bataillon, Guillaume P. Ramstein

**Affiliations:** Center for Quantitative Genetics and Genomics, Aarhus University, Aarhus C, 8000, Denmark; Bioinformatics Research Centre, Aarhus University, Aarhus C, 8000, Denmark; Peking University Institute of Advanced Agricultural Sciences, Shandong Laboratory of Advanced Agriculture Sciences in Weifang, Weifang, Shandong 261325, China; Plant Biology, Cold Spring Harbor Laboratory, Cold Spring Harbor, NY 11768, USA

**Author notes:** **Corresponding Author:** Natasha Johansen.

## Abstract

Quantitative genetic approaches such as genome-wide association studies and genomic prediction are widely used to identify favourable genetic variation, but they have limited resolution due to linkage disequilibrium. Comparative genomics approaches, especially Protein Language Models (PLMs), have emerged as powerful alternatives, by detecting phylogenetic residue conservation (PRC) across evolutionary time scales. However, the extent to which these tools can guide the detection of impactful variants for field agronomic traits is still unclear.

In this study, we used the pre-trained PLM ESM2 to predict PRC scores of nonsynonymous mutations segregating within a diverse panel of 387 accessions in sorghum (SAP). The distribution of fitness effects (DFE) of the same set of nonsynonymous mutations was inferred using unfolded site frequency spectra to assess whether the DFE distribution covaried with PRC scores. Furthermore, we estimated the load of putatively nonneutral mutations of SAP accessions and evaluated associations between this mutation load and phenotypic performance across multiple agronomic traits. Our results show that ESM2 can detect mutations associated with fitness-enhancing effects in SAP, as indicated by enrichments in positive selection signatures among the variants with positive PRC scores. Significant associations were also detected between phenotypic performance and mutation load for several agronomic traits, indicating that PLMs can identify functionally important genetic variation. However, these signals were not consistent across all traits in the SAP population. Altogether, our findings suggest that large language models may support breeding efforts, as PLM predictions covaried with fitness effects and captured agronomic performance for some traits in plant populations.

## 1. Introduction

During domestication, many modern crops underwent intensive artificial selection. While it ensured the rapid fixation of alleles favorable in domesticated conditions, population bottlenecks also increased the burden of deleterious variants (Beissinger et al. 2016; Liu et al. 2017; Moyers et al. 2018), especially in low recombination regions (Renaut and Rieseberg 2015). This extra burden of deleterious alleles, sometimes referred to as the cost of domestication can negatively affects agronomic performance (Valluru et al. 2019). To mitigate this issue, beneficial variation can be introduced or reintroduced through mutagenesis (Jiao et al. 2016, 2023), genome editing techniques, e.g., CRISPR (Barrangou and Doudna 2016; Rodríguez-Leal et al. 2017; Zhang et al. 2020), or by introgression of less advanced breeding material, including landraces (Meseka et al. 2013; Gorjanc et al. 2016) or wild relatives (Ananda et al. 2020). To effectively harness genetic variation, breeding efforts will benefit from tools that can prioritize variants that are most likely to be beneficial.

Methods used for this purpose include traditional approaches such as Genome-Wide Association Studies (GWAS) and Genomic Prediction (GP) (Meuwissen et al. 2001). GP models are routinely used to estimate the genetic performance (breeding values) of selection candidates based on genome-wide genetic information but are not aimed at identifying beneficial variants at high resolution, e.g., specific point mutations. GWAS can in principle detect alleles associated with phenotypic performance, i.e., quantitative trait loci (QTL) (Morris et al. 2013; Boyles et al. 2016). However, both GP and GWAS are affected by linkage disequilibrium (LD), where alleles at different loci are non-randomly associated within a population. Consequently, associations identified with GWAS may not pinpoint the causal variant, but rather noncausal variants in linkage with them (Flint-Garcia et al. 2003). The resolution of phenotype-genotype associations detected with GWAS is therefore dependent on the extent of LD in the population. Additionally, GWAS cannot easily discriminate between variants with unconditional effects, i.e. variants that are beneficial across different environmental/genetic contexts, and variants with conditional effects, i.e. only beneficial under certain environmental conditions.

In contrast, comparative genomic approaches aim at detecting phylogenetic conservation (PC) of nucleotides or amino-acid residues (PRC). Levels of observed PC are consistent with patterns of purifying selection across species and environments arising from functional constraints (Camps et al. 2007). The variation in the degree of PC primarily reflects the strength of purifying selection across the genome because genomic regions differ in functional importance. For example, mutations disrupting protein function or protein structure (Choi et al. 2012; Echave et al. 2016) or negatively impacting gene expression (Kremling et al. 2018) are expected to be eliminated through negative selection,.

Unlike GWAS, comparative genomics methods can identify functionally important regions at high resolution, down to specific base-pairs or amino-acid residues. Comparative Genomic approaches used for PC detection, can be based on alignment-based approaches which require multiple-species-alignment (MSA), but more recent deep learning methods do not require MSA (Sendrowski et al. 2025). Common MSA-based tools include SIFT (Ng and Henikoff 2001, 2003; Vaser et al. 2016) and GERP (Davydov et al. 2010). An important limitation of MSA-based approaches is that site conservation can only be assessed for genomic regions that can be aligned across sequences, hence variant effect predictions are not available for genomic regions lacking homologous sequence alignment. In contrast, advanced deep learning models, including biological language models are not constrained by sequence alignability. These include Protein Language Models (PLMs), such as the evolutionary scale model (ESM) (Rives et al. 2021), which can predict the effects of amino-acid substitutions on protein function. Importantly, the PLMs can generalize across proteins by capturing effects of mutations conditional on protein sequences.

Comparative genomics approaches, including PLMs, can predict the effects of individual variants by detecting selection signatures (Sendrowski et al. 2025). Among those, signatures of positive selection point to beneficial mutations, which include both adaptive mutations and restorative back mutations. Adaptive mutations are background-dependent (i.e., specific to certain genomic and environmental conditions), whereas back mutations have by definition background-independent effects (Charlesworth and Eyre-Walker 2007). This distinction is crucial, because back mutations may be detected through cross-species conservation, as captured by PLMs, whereas adaptive mutations can only be captured through clade-specific signals (Latrille et al. 2023, 2024).

Previous studies have investigated the association between phenotypic performance and the load of non-neutral variants for a range of crops, including sorghum, maize and potato (Yang et al. 2017; Valluru et al. 2019; Wu et al. 2023), barley and soybean (Kono et al. 2016). Most of these studies have, however, focused on deleterious mutations and relied on MSA-based approaches for variant effect prediction. In contrast to previous studies, we leveraged variant effect predictions from an advanced PLM (ESM2) and used these predictions to partition genetic variants into mutation classes including both deleterious and beneficial variant effects. This approach allowed us to efficiently validate the ability of PLM predictions to capture fitness effects and phenotypic performance within species.

The aim of our study is to leverage the pre-trained PLM ESM2 to detect putatively beneficial and deleterious variants with unconditional effects, i.e. genetic variants whose effects are consistent across long evolutionary time scales; and validate the average effect of putatively nonneutral variants on fitness and agronomic traits, using the Sorghum Association Panel (SAP) as a representative sample of species-wide diversity in plants (Boatwright et al. 2022).

## 2. Methods & Materials

This study combines two complementary parts: (i) **population genetics** analyses, including unfolded site frequency spectra (uSFS) and inference of the distribution of fitness effects (DFE); and (ii) **quantitative genetics** analyses, including genomic prediction (GP) models to assess the contributions of functionally prioritized variants to phenotypic performance.

### 2.1 Phenotypes and genotypes

The SAP diversity panel consisted of 400 accessions, of which 387 had been phenotyped. Agronomic traits investigated included quality traits (Amylose, Fat, Starch and Protein content), physiological traits (Panicle Length, Flag Leaf Height and Terminal Branch Length), production traits (Grain Number, Grain Weight and Grain Yield), and phenology traits (Days to Anthesis).Whole-genome sequence (WGS) data were derived from (Boatwright et al. 2022), while phenotypes were obtained from previous publications on the SAP (Boyles et al. 2016, 2017; Sapkota et al. 2020b, a).

### 2.2 Protein language model to detect nonneutral variants

Putatively non-neutral variants were identified using the PLM ESM2, specifically the esm2_t36_3B_UR50D sub-model (Lin et al. 2023). The model detects sites under purifying selection by analyzing sequence variation across diverse species, leveraging large-scale protein sequence data from the UniRef database (Suzek et al. 2007). For a given amino acid position in the protein sequence, the PLM estimates the probability of observing an alternative amino acid residue (*a*_*ALT*_) relative to the reference residue (*a*_*REF*_). This evolutionary score is calculated as the log-likelihood ratio 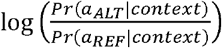, where Pr(*a*_*ALT*_) and Pr(*a*_*REF*_) are the predicted probabilities of the alternative residue and reference residue occurring at the specific position in the sequence, respectively, *context* refers to the sequence of the protein carrying the variant. In this study, predictions were derived from the canonical isoforms of the *Sorghum Bicolor* **BTx623** reference genome (v3), using Ensembl Plant release 55 annotations. PLM predictions were generated with the publicly available scripts from (https://github.com/ntranoslab/esm-variants).

To compare ESM2 with MSA-based approaches in analyses of allele frequency, we retrieved SIFT scores from the Ensembl Variant Effect Prediction database (McLaren et al. 2016), under the Ensembl Plants release In each protein isoform, SIFT scores between 0 and 1 quantify the PRC at each site by estimating the frequency of amino-acid residues across an MSA (Ng and Henikoff 2001). Similarly to ESM scores, SIFT scores are expected to be positively correlated with fitness.

### 2.3 Inference of the distribution of fitness effects in the population

The PLM predictions, or evolutionary scores, provide prior information regarding the potential fitness effects of variants. The unfolded Site-Frequency-Spectrum (uSFS) allows inference of the Distribution of Fitness Effects (DFE), which describes the probability distribution of scaled selection coefficients for the genetic variants of interest. The DFE can be used to assess whether sets of putatively beneficial variants, as predicted by the PLM, are collectively more likely to be beneficial based on the allele frequencies at which they segregate in the population. The uSFS summarizes the counts of derived alleles (*Der*) at different allele frequencies. This analysis thus requires that the ancestral allele (*Anc*) is known. Ancestral alleles were identified for all segregating polymorphic sites located in protein-coding regions alignable to two outgroups: maize (*Zea mays*) (Hufford et al. 2012) and a diploid wild relative of sugarcane (*Erianthus rufipilus*) (Wang et al. 2023).

To obtain the uSFS, two spectra are required: the neutral and the selected spectrum. The neutral spectrum accounts for demographic history and nuisance parameters, including genetic drift, while the selected spectrum contains the variants of interest. These spectra are represented by count vectors denoted by *p*_*N*_ and *p*_*S*_ respectively. Synonymous mutations at 4-fold degenerate sites (P_4_) were assumed neutral and were used to obtain *p*_*N*_, while nonsynonymous mutations at polymorphic 0-fold degenerate sites (P_0_) were used to obtain *p*_*S*_. Approximately 40,000 polymorphic sites, specially 0-fold and 4-fold degenerate sites, where alignable to outgroups. To obtain *p*_*S*_ for different mutation categories, the P_0_ sites are partitioned into ten equal-sized spectra, conditioned on the evolutionary score of the derived allele. The evolutionary scores are polarized to reflect the probability of *Der* relative to *Anc*, and are defined as:

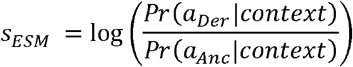

The spectrum of a mutation category *z*, denoted as 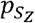, is then obtained from sites whose evolutionary scores *S*_*ESM*_ falls within the interval *I*_*z*_ = [*T*_*L,z*_,*T*_*U,z*_), where *I*_*z*_ represents the *Z*^*th*^ (bin) interval in the 10-partition. The set of sites used to obtain 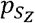 may be expressed as:

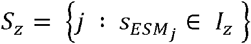

To infer a reliable DFE, it is necessary to account for variability in the number of mutational opportunities. This parameter will vary both across mutation categories, defined by the partition of evolutionary scores, and between the selected and neutral spectra, 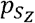 and 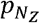. For a given spectrum, let *L*_*S*_ represent the number of mutational opportunities for 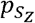, and *L*_*N*_ for 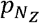. To get *L*_*s*_, we first estimate the proportion of *potential* mutations whose evolutionary score *S*_*ESM*_ falls within the interval *I*_*z*_. Specifically, for all 0-fold monomorphic sites, we determine the average evolutionary score of the possible point mutations that could occur at these sites. Let the number of monomorphic sites with evolutionary scores within the interval *I*_*z*_ be denoted by *M*_0,*z*_. Then *L*_*S*_ and *L*_*N*_ is estimated as:

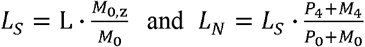

where *L* denote the total number of mutational opportunities across selected sites, *M*_0_ (*M*_0,*z*_) is the count of all (prioritized) 0-fold monomorphic sites, and *M*_4_ is the count of 4-fold monomorphic sites in regions alignable to outgroups. The DFE was fitted separately for each spectrum 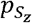, and selection coefficients *S* were scaled by the effective population size: *S* = 4*N*_*e*_*s*. The DFEs are modeled as a two-component mixture: a reflected gamma distribution for deleterious mutations (*s* < 0) with shape parameter b and mean *S*_*d*_, and an exponential distribution for beneficial mutations (*s* ≥ 0) with mean *S*_*b*_, and the parameter *P*_*b*_ representing the proportion of beneficial mutations, i.e., the mixture weight.

### 2.4 Linkage disequilibrium analysis

The decay of LD was estimated through the pairwise squared Pearson correlation (r^2^) as a function of physical distance, adjusted for population structure and kinship following the approach in (Mangin et al. 2012) and implemented in (Skovbjerg et al. 2025). We examined whether the rate of LD decay and baseline LD differed among predefined SNP categories, i.e., sites partitioned into ten mutation categories based on the evolutionary score for *Der* at each site. LD decay was modelled by fitting a generalized linear model (GLM) with a Gamma error distribution and inverse link function. Nested models were compared using likelihood ratio tests to determine whether (i) baseline and background LD differed between mutation categories (intercept) and (ii) rate of LD decay varied significantly between mutation categories (slope). Sites with a minor allele frequency (MAF) below 0.05 were excluded from this analysis.

### 2.5 Weighted mutation load

For all accessions we estimate the individual weighted mutation load which is given as:

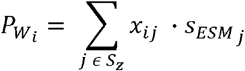

Where *x*_*ij*_ is the count of derived alleles in accession *i* at site *j*, and 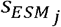 is the evolutionary score of the derived allele. The weighted mutation load, 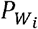 is therefore a measure of whether a given individual is enriched for putatively nonneutral alleles.

### 2.6 Genomic Prediction models

Two complementary GP analyses were performed:

i. **Mean partition:** We investigated whether there is a relationship between phenotypic performance and the load of putatively nonneutral mutations in each mutation category.
ii. **Variance partition:** We tested whether the distribution of variant effects differed across mutation categories.

In both analyses, sites were partitioned according to the same ten mutation categories defined in the previous section (2.3-2.4). Consequently, variant prioritization was restricted to sites located in protein-coding regions that are alignable to outgroups.

#### 2.6.1 Genomic relationships matrices

The genomic relationship matrix (GRM), **G**, was computed following (VanRaden 2008):

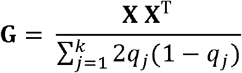

where **X** is the centered genotype matrix. The scaling factor 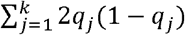 is the expected heterozygosity at each site under Hardy-Weinberg equilibrium, *q*_*j*_ is the frequency of the alternative allele at site *j*.

#### 2.6.2 Baseline GP model

The baseline model is the Genomic Best Linear Unbiased Prediction (GBLUP) model (Habier et al. 2013):

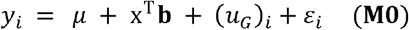

where *y*_*i*_ is the phenotypic performance for accession *i, µ* is the grand mean, and **x**^T^b a vector of fixed effects. The fixed effects include the first three principal components (PCs), from a principal component analysis (PCA) performed on the centered genotype matrix **X**, as well as the genomewide count of derived alleles across all 0-fold sites. These covariates account for population structure and the effect of being enriched for derived alleles, respectively. The random additive genetic effects are polygenic effects 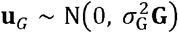 and residual errors 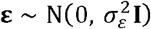, **I** being the identity matrix.

#### 2.6.3 Mean partition

The baseline model was extended by including mutation load as a fixed effect, defined as the total number of derived alleles across prioritized sites. For each accession *i*, mutation load was computed as the sum of derived alleles at sites where the evolutionary score of the derived allele falls within a predefined interval. The mutation load for accession *i*, within interval ***I***_***z***_ is defined as:

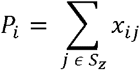

where *S*_*z*_ is a mutation category as defined above, *x*_*ij*_ is the count of derived alleles in accession *i* at site *j*. The term was included in the model as a fixed effect with coefficient *β*_*P*_.:

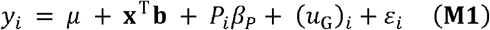

Wald tests and permutation tests were used to assess the statistical significance of the inferred effects of the prioritized variants on mean phenotypic performance 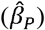.

#### 2.6.4 Variance partition

We extended the baseline GBLUP model (M0) by assuming that prioritized variants follow a different variant effect distribution than the baseline additive genetic distribution 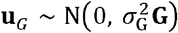. In the extended model, we partitioned the genetic variance into two components, **G** and **G**_P_, where the variance component for **G**_P_ represent the additive genetic variation attributable to genetic differences at the prioritized sites. We therefore tested whether the variance in variant effects at prioritized sites included in **G**_P_ was significantly different when compared to genome-wide variants included in **G**. To construct **G**_P_, we used the genotype matrix **X**_z_ corresponding to the set of prioritized sites *S*_*z*_.

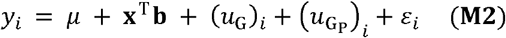

where 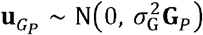.

Log-Likelihood Ratio (LLR) tests were performed based on Restricted maximum likelihoods (REML) to compare nested random models. Likelihood ratios were calculated as: □□ = −*2* [□_*0*_(□) − □_*1*_(□)], where □_*1*_(□) is the log-likelihood of the tested model, and □_*0*_(□) the log-likelihood of the null model. Under the null hypothesis, likelihood ratios follow a chi-square distribution, where the degrees of freedom equal the difference in number of fitted model parameters.

### 2.7 Model comparison and prediction ability

To evaluate the performance of GP models, a leave-one-genetic-cluster-out validation scheme was used, where the accessions were assigned into six genetic clusters based on genetic relatedness, following a previous SAP publication (Boatwright et al. 2022). For each iteration, all accessions in one cluster were completely held out as a test set, and model was trained on the remaining accessions. This was repeated for the remaining clusters. The prediction accuracy was determined for each cluster, as the Pearson correlation between predicted and observed phenotypic records 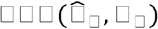. Finally, models were evaluated based on the mean prediction ability across genetic clusters, hereafter PA.

To assess the statistical significance of PA, a total of *n* = 5,000 permutations were performed with empirical p-values given as P = (1 + r)/ (1 + n), where n is the total number of permutations, and *r* the number of permutations with effects that is equal or more extreme than the observed effect (North et al. 2002).

### 2.8 Software

Linear-Mixed-Models (LMM) were fitted with the R packages MM4LMM version 3.0.2 (Laporte et al. 2022) and qgg (Rohde et al. 2020). The R package FastDFE was used to generate the uSFS and estimate their associated DFEs (Sendrowski and Bataillon 2024). All analyses were run in R 4.2.0 (Development Core Team) within the high-performance computing cluster GenomeDK.

## 3. Results

### 3.1 Evolutionary scores correlate with allele frequency in diversity panel

The SAP displayed moderate population structure, with the first two PCs explaining about 15 % of the genomic variation in the panel. The weighted mutation load differed significantly among genetic clusters (one-way ANOVA, F = 65.7, p < 0.001), indicating that certain genetic clusters are enriched for putatively beneficial mutations. Tukey’s HSD post hoc tests indicated that genetic clusters 2 and 3 exhibit significantly higher weighted allele loads than cluster 1 and clusters 4-6, suggesting enrichment of alleles predicted to be beneficial in these genetic clusters (**Fig. 2A**). Beneficial variants are expected to be under positive selection and therefore occur at higher allele frequencies than deleterious variants. Evolutionary scores derived from either SIFT (scores derived from MSA-based approach, used here as a baseline) or ESM (scores derived from a PLM-based approach) are positively associated with allele frequency in the SAP, suggesting a positive association with fitness effects (**Fig. 2B**). For both types of evolutionary scores, we found a significant association between the frequency of alternate alleles and the evolutionary score (ESM: p < 0.0001, Adjusted R^2^ = 0.060) and (SIFT: p < 0.0001, Adjusted R^2^ = 0.045) based on a generalized additive model, with chromosome number included as a categorical covariate (Wood 2017).

**Figure 1:**
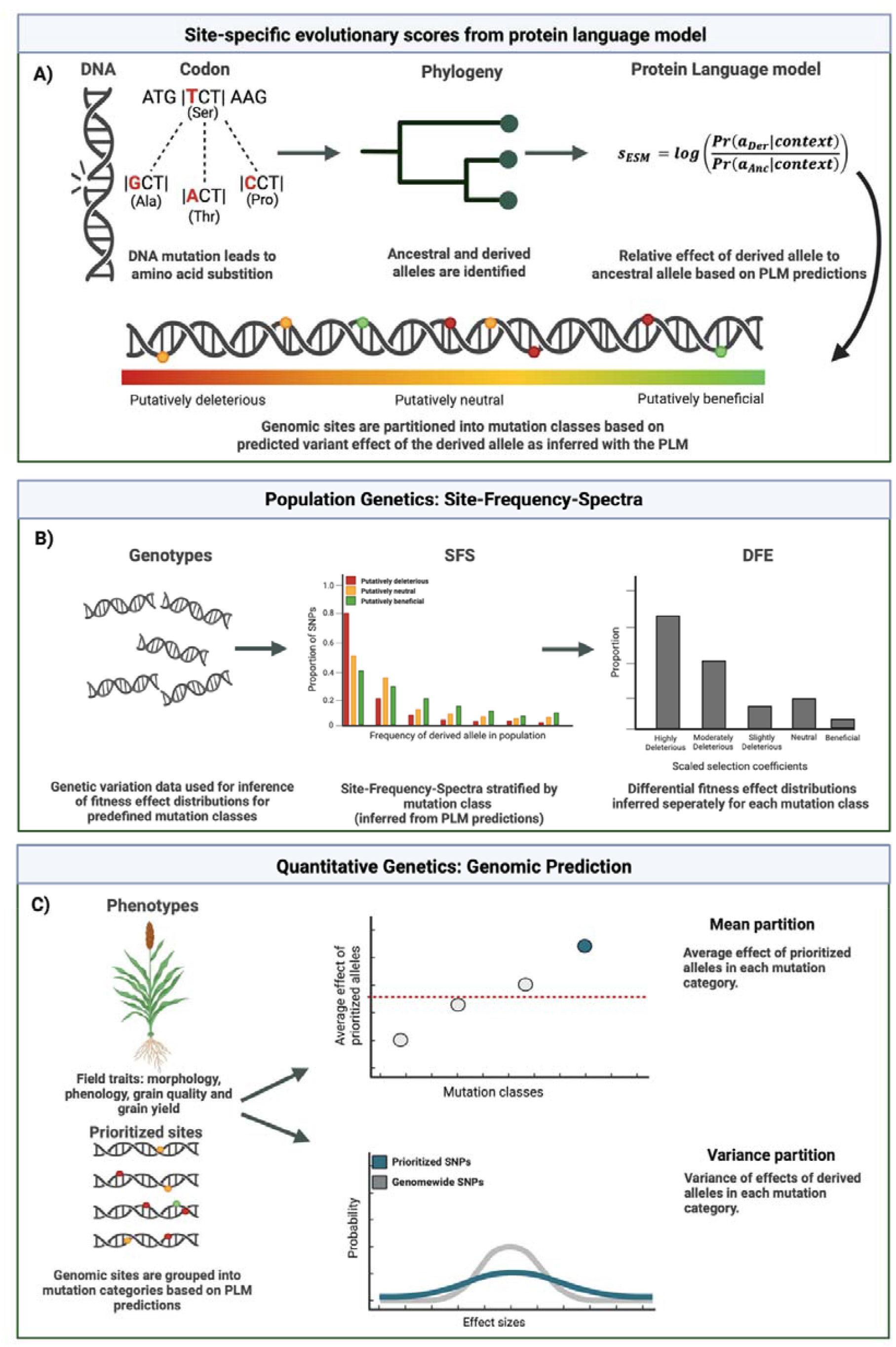
Overview of the analyses conducted in this study. Created in BioRender. Johansen, N. (2026) https://BioRender.com/8eg4qee

**Figure 2:**
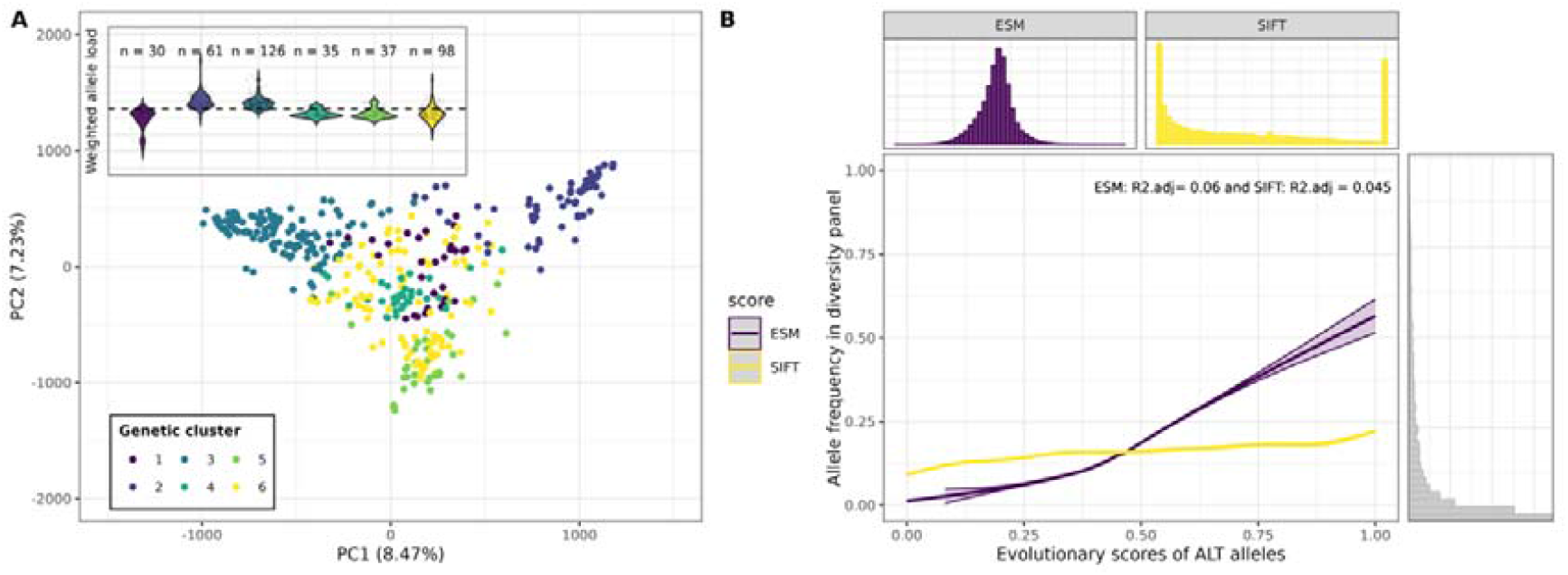
PLM-based evolutionary scores show stronger correlation with allele frequency than MSA-based scores. **(A)** Principal component analysis (PCA) in the Sorghum Association Panel (SAP) based on genome-wide SNP data. Points represent individual accessions, with colors indicating the assigned genetic cluster. The embedded violin plot shows the distribution of weighted mutation load, where the weights correspond to evolutionary scores of the derived allele (PLM-based), normalized to a range between –1 and 1. An increase in weighted mutation load reflects enrichment for mutations predicted to have fitness-enhancing effects. The stippled line indicates the average weighted mutation load across all accessions, sample size for each genetic cluster is shown above the distributions. (**B**) Frequency of the ALT alleles in the SAP, as a function of min-max-normalized evolutionary scores, polarized to reflect the probability of the ALT allele relative to REF, including all missense variants in protein-coding regions. Associations are shown for two evolutionary score metrics, ESM (PLM-based) and SIFT (MSA-based). The distribution of ALT allele frequencies shown in the right-hand panel, while the distribution of the two evolutionary scores is shown in the upper panel.

The stronger association achieved with ESM suggests that these PLM-derived scores may better predict fitness effects in the SAP, compared to MSA-based scoring approaches. Importantly, ESM scores allowed for a finer partition of variants, due to their continuous distribution, contrary to SIFT scores, whose values were concentrated at exactly 0 or 1 (**Fig. 2B**).

### 3.2 The ESM protein language detects fitness-enhancing variants in SAP

To investigate the relationship between evolutionary scores from ESM2 and fitness effects, we conducted analyses of LD decay and fitness effect distribution in the SAP. The LD decay analysis indicated that both background levels of LD (χ^2^-test, p < 0.001) and the rate of LD decay (χ^2^-test, p < 0.001) varied significantly between different mutation categories. These differences may reflect differences due to selection, i.e., background selection for putatively deleterious mutations and positive selection for putatively beneficial mutations. Differences in LD decay between mutation categories suggested lower haplotype diversity around putatively beneficial variants, as expected under selective sweeps (**Fig. 3B)**.

**Figure 3:**
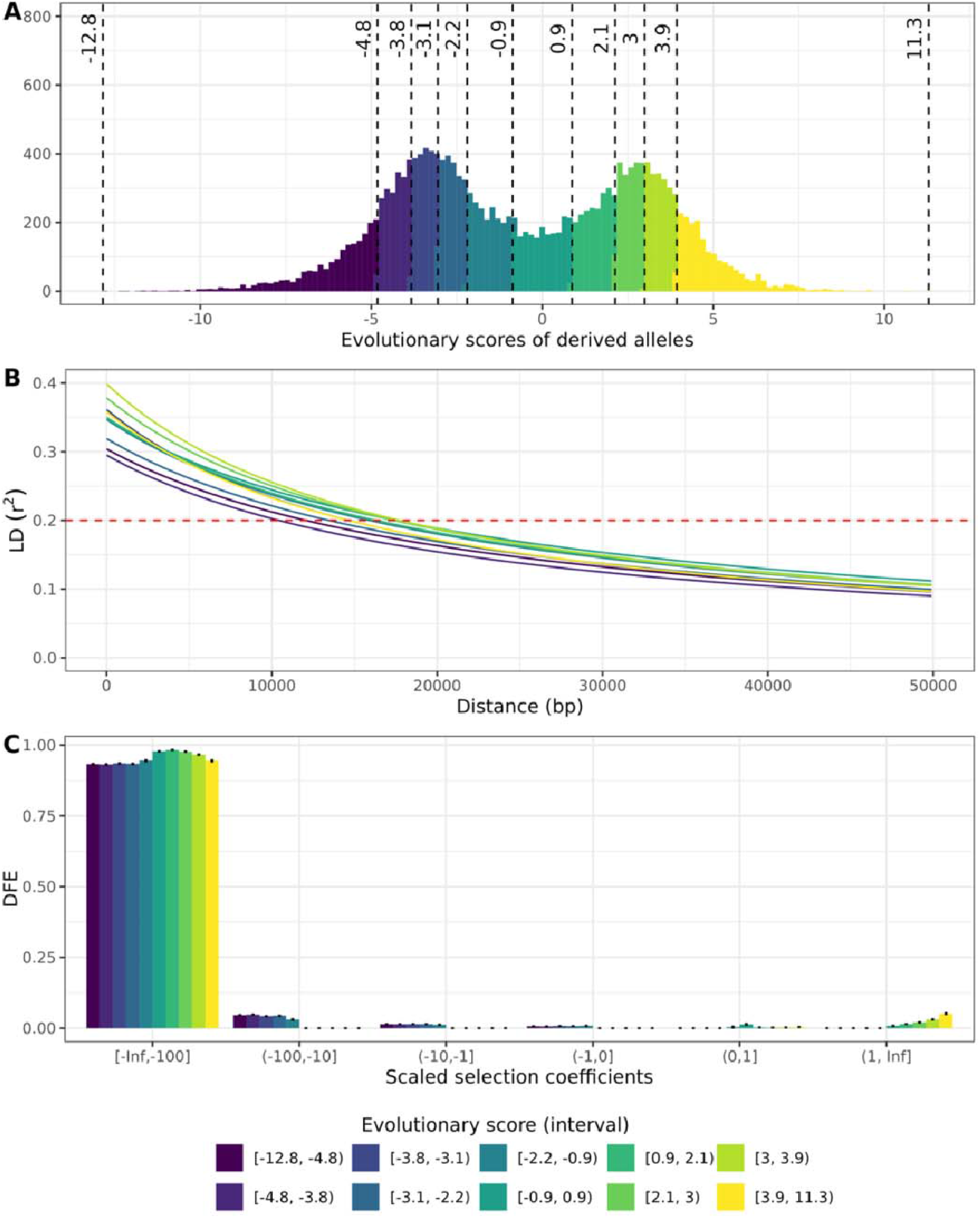
Rate of LD decay and the distribution of fitness effects across mutations categories defined by evolutionary scores. **(A)** Genome-wide distribution of evolutionary scores predicted by the Protein-Language-model (PLM) ESM2. Scores were generated for all 0-fold degenerate sites in regions alignable to outgroups. Scores are polarized to reflect the (log) ratio probability of observing the derived allele (Der) relative to the ancestral allele (Anc). Dashed lines indicate the boundaries separating deciles of evolutionary scores for the ten mutation categories used to generate the spectra. (**B**) Rate of LD decay (r^2^) as a function of genomic distance (bp), shown for each mutation category, with the red stippled line indicating the 0.2 threshold. (**C**) Distribution of Fitness Effects (DFE) distributions, inferred for each of the ten mutation categories, shown with 95% confidence intervals (based on 100 bootstrap replicates).

If within-species fitness effects covary with PLM-based predictions, mutations with *s*_*ESM*_ >0 will segregate at higher allele frequencies than neutral mutations. Specifically, if a spectrum 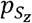 is derived from sites where mutations are predicted to be beneficial, the corresponding DFE should likewise be shifted toward positive selection coefficients (4*N*_*e*_ *s* > 0).

Conversely, if 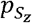 is derived from sites where the mutations have *s*_*ESM*_ < 0, these variants are expected to segregate at low frequencies due to negative selection and the DFE should thus be shifted downward toward negative selection coefficients (4*N*_*e*_ *s* < 0).

Our results show that all spectra are characterized by DFE distributions shifted toward highly deleterious mutations (4*N*_*e*_ *s* « −1 *or* − 10), indicating that the proportion of highly deleterious mutations is not reduced in the spectra obtained from putatively beneficial mutations (**Fig. 3C**; **Fig. S1**). However, the proportion of beneficial mutations increased markedly for mutation categories consisting of putatively beneficial mutations. Specifically, the proportion of beneficial mutations (1 < 4*N*_*e*_ *s* ≤∞) increased consistently from 0% for the spectra obtained from putatively deleterious or neutral mutations to 6% for the 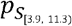 spectrum, i.e., the spectrum obtained from mutations predicted to be most beneficial by the PLM (**Table 1**).

**Table 1:** Maximum likelihood analysis for fitness effects (uSFS/DFE). Model parameters shown for the gamma model.

| ESM score interval | LLR | Gamma-model |  |  | Sd | Sb | pb |
| --- | --- | --- | --- | --- | --- | --- | --- |
| | | ML $\alpha$ | ML b | ML $\theta$ | | | |
| [-12.8, -4.8) | -202.3 | 0 | 0.51 | 0.0 | -13,217 | $0.10 \cdot 10^{-4}$ | 0 |
| [-4.8, -3.8) | -216.3 | 0 | 0.52 | 0.0 | -11,005 | $0.30 \cdot 10^{-4}$ | 0 |
| [-3.8, -3.1) | -202.4 | 0 | 0.46 | 0.0 | -22,139 | $0.10 \cdot 10^{-3}$ | 0 |
| [-3.1, -2.2) | -157.2 | 0 | 0.47 | 0.0 | -17,572 | $0.20 \cdot 10^{-4}$ | 0 |
| [-2.2, -0.9) | -85.7 | 0.28 | 0.39 | 0.0 | -100,000 | $0.10 \cdot 10^{-4}$ | 0 |
| [-0.9, 0.9) | -101.2 | 0.99 | 10.0 | 0.0 | -100,000 | 1.0 | 0.004 |
| [0.9, 2.1) | -260.8 | 1 | 10.0 | 0.0 | -100,000 | 5.8 | 0.02 |
| [2.1, 3) | -413.1 | 1 | 10.0 | 0.0 | -100,000 | 9.7 | 0.02 |
| [3, 3.9) | -449.3 | 1 | 10.0 | 0.0 | -100,000 | 12.1 | 0.03 |
| [3.9, 11.3) | -412.7 | 1 | 10.0 | 0.0 | -100,000 | 13.3 | 0.06 |

\* ML $\alpha$ : Maximum likelihood estimate of the shape parameter of the gamma distribution of fitness effects.
\* ML b: Maximum likelihood estimate of the scale parameter of the gamma distribution; larger values indicate the DFE, on average, is skewed toward nonneutral effects.
\*ML $\theta$ : The ancestral allele misspecification rate, i.e., the proportion of alleles where the derived allele is incorrectly identified.

Thus, our analyses suggested that evolutionary scores covary with within-species fitness effects. Given that the proportion of highly deleterious mutations remained constant or even increased for the spectra derived from putatively beneficial mutations, it appeared that ESM-based prioritizations include false positives. Nevertheless, given the strong association with allele frequency (**Fig. 2B**), the observed trends in LD decay (**Fig. 3B**), and the consistent increase in the probability of beneficial mutations across ESM intervals (**Fig. 3C; Table 1**), our result provide evidence for a significant enrichment in beneficial effects among variants prioritized by high PLM-based scores.

### 3.3 Effects of ESM-prioritized variants may improve genomic prediction of fitness-related traits

We conducted quantitative genetics analyses to determine the relationship between evolutionary scores and the mean or variance of variant effects for agronomic traits in the SAP (respectively, mean and variance partition). In mean partition analyses, significant associations between phenotypic performance and the mutation load were generally observed for morphological traits, including Flag Leaf Height, Panicle Length and Terminal Branch Length (**Fig. 4A, Table 2**). These associations were mostly observed when prioritization was based on sites with the most extreme evolutionary scores. Specifically, the variants with very low ESM scores in the interval was found to be, on average, positively associated to Flag Leaf Height, Panicle Length (Fig 3A) and Terminal Branch Length (Fig. S3).

**Table 2:**
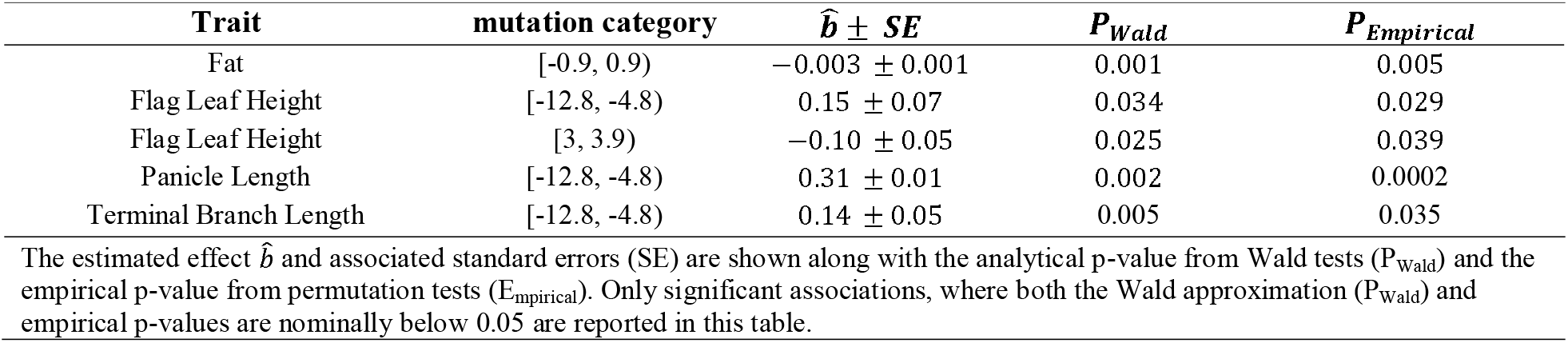
Estimated average effect of prioritized alleles for the different SNP categories.

| Trait | mutation category | $\hat{b} \pm SE$ | $P_{Wald}$ | $P_{Empirical}$ |
| --- | --- | --- | --- | --- |
| Fat | [-0.9, 0.9) | $-0.003 \pm 0.001$ | 0.001 | 0.005 |
| Flag Leaf Height | [-12.8, -4.8) | $0.15 \pm 0.07$ | 0.034 | 0.029 |
| Flag Leaf Height | [3, 3.9) | $-0.10 \pm 0.05$ | 0.025 | 0.039 |
| Panicle Length | [-12.8, -4.8) | $0.31 \pm 0.01$ | 0.002 | 0.0002 |
| Terminal Branch Length | [-12.8, -4.8) | $0.14 \pm 0.05$ | 0.005 | 0.035 |

The estimated effect $\hat{b}$ and associated standard errors (SE) are shown along with the analytical p-value from Wald tests ( $P_{Wald}$ ) and the empirical p-value from permutation tests ( $P_{Empirical}$ ). Only significant associations, where both the Wald approximation ( $P_{Wald}$ ) and empirical p-values are nominally below 0.05 are reported in this table.

**Figure 4:**
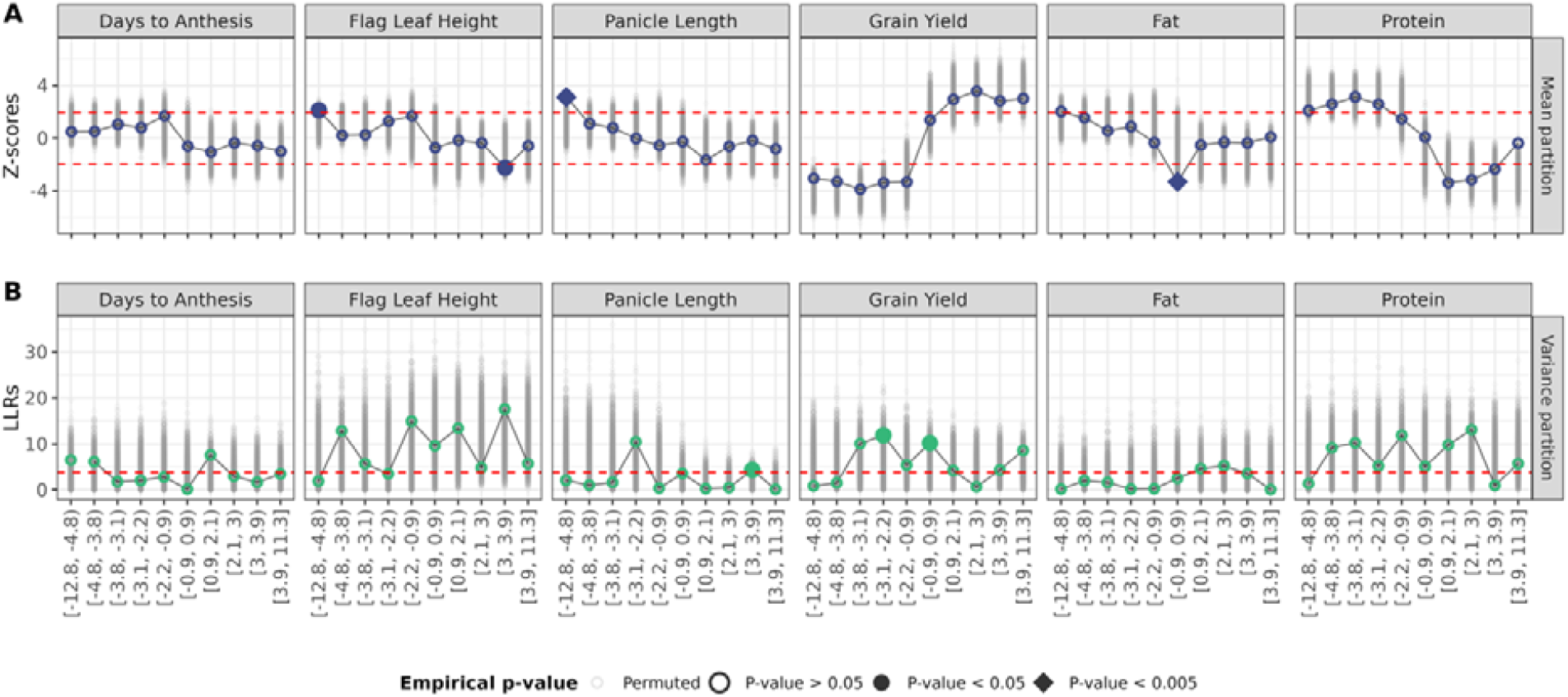
Impact of ESM-based prioritizations on genomic prediction model performance. **(A)** Mean partition: **Mutation** categories given as intervals of evolutionary scores for the derived allele shown on the x-axis. Points represent the estimated average effect of the prioritized alleles, i.e., alleles contributing to mutation load, standardized as Z-scores (estimate divided by its standard error). Red dashed lines indicate the threshold for significance at the 5% level based on Wald tests. **(B)** Variance partition: Differences in log-likelihood ratios () between the baseline model M0 and extended model M2 are shown. Red dashed lines indicate threshold for significance at the 5% level based on LLR tests. For both plots, the empirical significance, calculated based on 5000 permutations, is indicated by the point shape.

Contrary to expectations, the strongest overall association was observed for the trait Fat (lipid content), where accessions enriched for mutations with evolutionary scores between, i.e., predicted as neutral by the PLM, exhibited decreased lipid content in the seeds. In variance partition analyses, differences in log-likelihood ratios between the baseline GP model (M0) and the extended GP model (M2) generally did not consistently indicate an improved model fit for the traits that showed significant associations in the mean partition analysis (**Fig. 4B, Table 3**). This suggests that the potential benefit of functional prioritization of variants based on evolutionary scores varies across traits and depends on the modelling approach (mean vs. variance partition). Because the ESM2 model captures evolutionary constraint across a broad spectrum of species and environments, traits which are weakly related to plant fitness or subject to differential selection across environments may show weak associations with the variants prioritized/ranked by ESM2. The benefit of utilizing phylogenetic conservations captured by ESM2, seems highly trait-dependent, possibly because some traits (e.g., anthesis date) are impacted by clade- or environment-specific effects that are missed by the PLM.

**Table 3:** Results from the variance partition analysis comparing a traditional genomic prediction model to an extended model incorporating functional constraint.

| Trait | Mutation category | LLR | $P_{LLR-test}$ | $P_{Empirical}$ |
| --- | --- | --- | --- | --- |
| Panicle Length | [3, 3.9) | 4.3 | 0.038 | 0.022 |
| Terminal Branch Length | [-3.1, -2.2) | 22.55 | < 0.001 | 0.042 |
| Terminal Branch Length | [3, 3.9) | 8.6 | 0.003 | 0.002 |
| Grain Number | [-3.1, -2.2) | 21.1 | < 0.001 | 0.01 |
| Grain Yield | [-3.1, -2.2) | 11.9 | 0.006 | 0.045 |
| Grain Yield | [-0.9, 0.9) | 10.2 | 0.001 | 0.048 |

The difference in Log-likelihood-ratios (LLRs) between the baseline model (M0) and the extended model (M2) are shown, along with the analytical p-value from LLR tests ( $P_{LLR-test}$ ) and the empirical p-value from permutation tests ( $P_{Empirical}$ ). Only significant associations, where both analytical empirical p-values are nominally below 0.05 are reported in this table.

Overall, the baseline GP model (M0) showed moderate prediction accuracy across traits, ranging from 0.14 for Amylose to 0.45 for Grain Number (**Table 4**). The extended models (M1 and M2) did not consistently improve PA across traits, compared to the baseline model M0 (**Fig. 5**). The largest improvements were observed for the traits Protein, Panicle Length and Grain Yield. Specifically, Grain Yield showed a 7% increase in prediction accuracy under the M2 model assuming a different effect distribution for derived alleles with very high ESM scores, in [3.9, 11.3). Panicle Length showed the largest improvement under the M2 model, which assumed differential variant effect distributions for sites where the derived allele had scores in the interval [-3.1, -2.2). For Protein, the greatest improvement (8%) occurred with M1, where we modeled the average effects of derived alleles having moderately high ESM scores, in [2.1, 3).

**Table 4:** Average prediction accuracy (PA), across genetic clusters, shown for the baseline model (M0).

| Trait | Prediction accuracy ± SE |
| --- | --- |
| Amylose | $0.14 \pm 0.08$ |
| Cal.g | $0.35 \pm 0.13$ |
| Days to Anthesis | $0.24 \pm 0.10$ |
| Fat | $0.27 \pm 0.06$ |
| Flag Leaf Height | $0.33 \pm 0.14$ |
| Grain Number | $0.45 \pm 0.04$ |
| Grain Weight | $0.31 \pm 0.09$ |
| Grain Yield | $0.34 \pm 0.06$ |
| Panicle Length | $0.32 \pm 0.11$ |
| Protein | $0.23 \pm 0.07$ |
| Starch | $0.41 \pm 0.09$ |
| Terminal Branch Length | $0.31 \pm 0.11$ |

**Figure 5:**
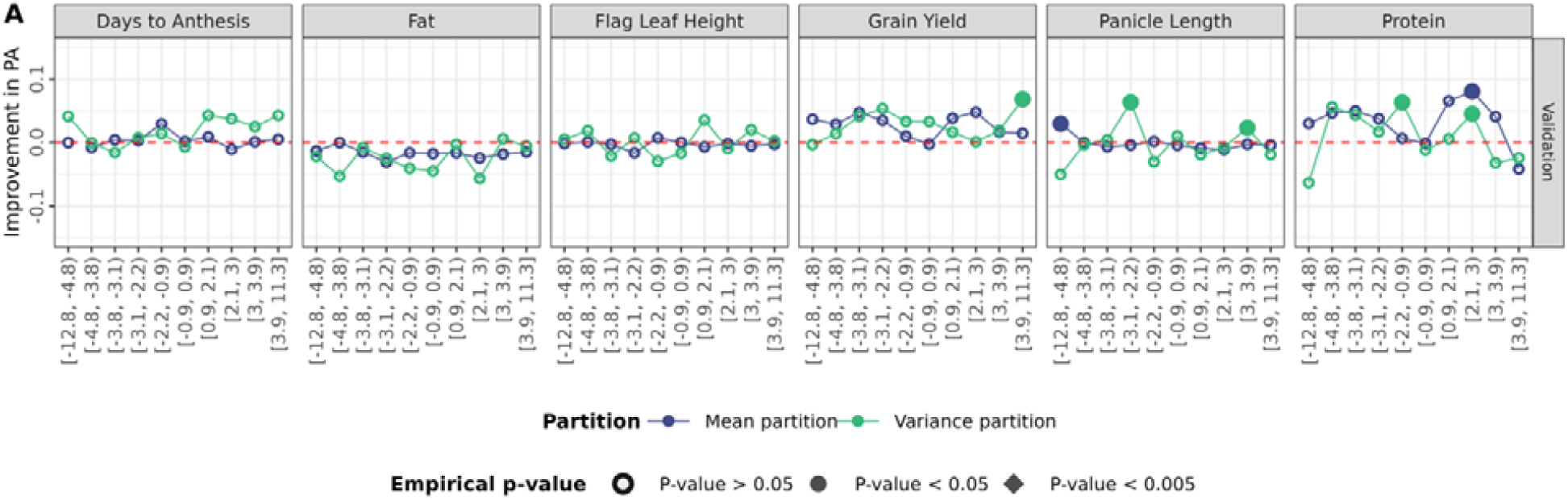
Functional prioritization of variants based on PLM variant effect predictions improves genomic prediction models’ ability to predict the genetic performance of novel germplasm. Improvement in predictive ability (PA) with the extended model M1 (mean partition) and M2 (Variance partition). Empirical significance derived from permutation tests are indicated by point shape.

These results underline that some traits benefit from evolutionary constraints informed GP models, while others do not. This is in accordance with our expectation, as we primarily expect certain fitness-related traits to benefit from this type of GP models. Improvements in PA were generally observed for specific combinations of traits and mutation categories, consistent with expectations that not all traits are impacted by the same mutation categories, due to differences in genetic architecture and biological effects across categories. Furthermore, it is also important to note that only sites with ancestral allele annotations were prioritized in either of the modelling approaches. These sites represent only a small fraction, approximately 10%, of the polymorphic sites within protein-coding regions. Given that the traits investigated in the study are quantitative traits, further gains in predictive ability may be achieved if the analysis is extended to a larger proportion of the genome. Finally, one challenge for detecting functional enrichment among putatively beneficial variants was collinearity with other variables in GP models. Especially with mean partition models, the collinearity of mutation loads and other covariates in the model (as quantified by the variance inflation score) was especially high with variants prioritized by positive ESM scores (**Fig. S4**).

## 4. Discussion

### 4.1 Detecting selection signatures with protein language models

Compared to previous studies on mutation loads in sorghum which used MSA-based approaches for identifying deleterious mutations (Valluru et al. 2019; Lozano et al. 2021), here we assessed the potential of a PLM to infer variant effects, based on both population genetics and quantitative genetics analyses. Our results support that PLM predictions are associated with variant effects. Specifically, we found that variants predicted to be beneficial by the PLM appear at higher allele frequencies within our focal population than expected under neutral processes alone, indicating that these genetic variants may be under positive selection in this population, presumably because they confer a fitness advantage (**Fig. 2B, Fig. 3C**). These findings are in agreement with previous studies based on SIFT scores (Chen et al. 2022; Latrille et al. 2024). Contrary to SIFT scores, whose distribution is strongly concentrated at extreme values 0 and 1, PLM scores are continuous values which allowed us to partition variants into several classes, distinguishing variants with ESM scores ranging from low to high while ensuring each class has the same size (**Fig. 3A**).

Despite a noticeable increase in the proportion of beneficial mutations with the putatively beneficial mutation categories, we also observe a high proportion of deleterious variants, even in categories with the highest evolutionary scores (**Fig. 3C**). This result suggests that additional information may refine the prioritization of variants beyond the information provided by ESM2. In the future, additional sources of information may be used to detect beneficial mutations with even higher accuracy. For example, information about protein expression, structure, and function may point to variants impacting fitness through mistranslation or disruption of essential biological function (Choi and Hannenhalli 2013; Zhang and Yang 2015). Further functional enrichment studies are needed to determine how such heterogeneous sources of information can be integrated to prioritize variants optimally.

### 4.2 The potential for protein language models for detecting variant effects on agronomic performance

Significant associations between phenotypic performance and mutation load were primarily observed for morphological traits, including Flag Leaf Height, Terminal Branch Length and Panicle Length, rather than production traits like grain number, grain weight, and grain yield. These associations were generally found between phenotypic performance and the mutation load of putatively deleterious alleles, particularly those predicted to be most deleterious (**Fig. 4, Fig. S3**), whereas significant associations between the load of putatively beneficial mutations and phenotypic performance were detected only for Flag Leaf Height. The weaker associations for production traits may result from their highly polygenic genetic architectures. In addition, mutation loads were calculated from ∼2,000 sites per mutation category, which may reduce power for traits influenced by many loci or by genes outside the regions with available ancestral allele annotations. Other studies using nucleotide phylogenetic constraint (PC) report that either observed or predicted nucleotide PC can be used for refinement of GP models when predicting grain yield in maize (Yang et al. 2017; Ramstein and Buckler 2022) Consistent with these studies, we achieved improvements in GP accuracy for grain yield when prioritizing variants with very high evolutionary scores (**Fig. 5**). However, these results should be replicated in other panels. In particular, future studies in breeding populations – rather than diverse panels like the SAP – should test whether scores derived from PLMs can effectively increase genetic gains in practical breeding programs.

### 4.3 Detecting beneficial alleles in plant breeding programs

In the context of plant breeding, we must also consider that natural selection and artificial selection may not align. Variants predicted to be deleterious based on evolutionary constraint may therefore be favored in breeding programs. This highlights the importance of understanding how enrichment of putatively beneficial alleles, as predicted by the ESM2, affects multiple traits simultaneously, and the direction of these effects. The current study utilizes an approach similar to the one presented in (Valluru et al. 2019), based on GERP and SIFT scores to identify putatively deleterious mutations in sorghum. They reported that the load of putatively deleterious variants was negatively correlated to phenotypic performance for a range of fitness related traits, i.e. plant height, dry biomass and tissue starch content (Valluru et al. 2019). Our results, suggests that the load of putatively deleterious variants was positively related to height-related traits (Flag Leaf Height and Panicle Length), which disagrees with their results (Valluru et al. 2019). These contrasting results may have resulted from differences in data and modelling approaches: in this study we investigated different populations and accounted for population structure to avoid confounding effects of demography. In general, we detected significant associations between phenotypic performance and mutation load under very stringent prioritizations, consistent with previous functional enrichment studies which show consistent increase in the magnitude of variant effects as variant prioritization grow more stringent (Ramstein and Buckler 2022; Zhai et al. 2025). One notable exception was lipid content (Fat), for which putatively neutral variants appeared to be functionally enriched. These results highlight the need to better understand for which traits implementation of PLM predictions for germplasm assessments may be advantageous. In particular, they suggest that variants with evolutionary scores close to zero may still be worth considering in breeding applications.

Given that variants with the greatest impact on field traits were generally found in mutation classes associated with the most extreme mutation categories, breeders should strive toward identifying and inducing the variants with the highest evolutionary scores in each plant genome and, equivalently, reverting the variants with the lowest evolutionary scores. However, these represent a relatively small pool of genetic variants, and the expected gains from such a narrow investment may be limited. Indeed, the assumption that few beneficial variants may contribute markedly to increasing genetic gain may be unrealistic (Khaipho-Burch et al. 2023). Therefore, an optimal breeding strategy may rely on the synergy between genomic selection, where estimation of genomic breeding values is guided by variant prioritization, and precision editing targeted at the most impactful variants. In genomic selection, predicted variant effects may be integrated through mean and/or variance partitioning approaches, as demonstrated in this study. Such models could be further extended to include all genomic variants, weighting them differentially according to their evolutionary scores to maximize predictive accuracy (Wu et al. 2023). In precision editing, evolutionary and mechanistic information about variant effects may be used to pinpoint the variants most likely to have beneficial impacts on traits of interest (Glaus et al. 2025).

### 4.4 Marker-based vs. haplotype-based approaches for detecting beneficial variation

Our study focused on scoring individual SNPs based on PLM predictions. This scoring approach is best suited to target individual variants for precision editing. In genomic selection, a promising extension of our approach is to model haplotype-based rather than SNP-based variation. Our approach did not account for the indirect effects of linked, non-prioritized variants that are inherited alongside prioritized alleles as part of a single haplotype. Thus, even if a prioritized variant independently confers a fitness advantage, the haplotype carrying the beneficial variant might be enriched for deleterious background mutations. In such cases, the net fitness association of the haplotype could be negative. Highly selfing species of plants like sorghum are particularly susceptible to this phenomenon, known as Hill-Robertson-interference (HRi) (Hill and Robertson 1966; Daigle and Johri 2024).

Furthermore, the extensive genetic diversity and population structure inherent in breeding germplasm may result in haplotypes that are unique to specific subpopulations. This heterogeneity can further impede the detection of significant associations between phenotypic performance and mutational load, as the magnitude and direction of HRi may vary significantly across different genetic backgrounds. To minimize this bias, studies of diverse germplasm may consider methods to model variation among haplotypes. These include studies in maize and wheat, which indicated that landraces carry beneficial haplotypes that have been lost or overlooked during modern breeding (Mayer et al. 2020; Cheng et al. 2024). In similar contexts, future studies may leverage recent sequence-based deep learning techniques to capture haplotype variation efficiently. For example, in maize, prediction of protein structure across gene haplotypes, based on AlphaFold2, has proven useful to explain and predict agronomic traits (Wang et al. 2026). Similar approaches with PLMs (e.g., based on variation in PLM sequence representations or mutation load across haplotypes) may facilitate detection of fitness-enhancing haplotypes found in underutilized material.

### 4.5 Conclusions

In this study, we conducted population genetics analyses, in which 0-fold degenerate sites were grouped into ten mutation categories based on evolutionary scores of their potential missense mutations, as estimated by ESM2. This approach allowed us to infer the underlying DFE, from which we determined that evolutionary scores covary with fitness effects (as measured by scaled selection coefficients).

We further employed quantitative genetic approaches, specifically GP models, to evaluate associations between phenotypic performance and mutation load across distinct mutation categories. Our findings indicate that PLM-based scores covary with realized within-species fitness effects, and that putatively nonneutral mutations exert a significant impact on field traits. Associations between phenotypic performance and mutation load were most significant for morphological traits, while no significant associations were observed for production traits (grain weight, grain number or grain yield). The potential advantage of accumulating putatively beneficial variants, as identified by the PLM ESM2 may therefore depend on the breeding goal for each population. In our GP analysis, we observed improvements in predictive ability when comparing a traditional GBLUP model to an extended GBLUP model where functional prioritization was performed based on PLM predictions. This suggests that traditional quantitative genetics approaches, including GP models, may benefit from integrating PLM-based functional prioritization of variants, highlighting their potential utility in breeding applications.

## Acknowledgements

We thank Doreen Ware (Cold Spring Harbour Laboratory) for her valuable guidance and support in the bioinformatics analyses, in addition to guidance and helpful discussions.

## Declarations

### Data and code availability statement

Genotypic and phenotypic data were obtained from (Sapkota et al. 2020b, a) and is publicly available. Evolutionary scores (ESM and SIFT scores) in addition to degeneracy and ancestral allele annotations are available at Zenodo https://doi.org/10.5281/zenodo.19494658. Genomic relationship matrices (GRMs) and genomic prediction inputs can be fully reproduced from the publicly available data using the provided scripts in the public repository.

The scripts used in this analysis are publicly available at https://github.com/TashaDear/BackOnTrack_sorghum.git. The scripts deposited in the repository include the workflows for calculation of mutation loads and GRM construction in addition to the unpermuted and permuted genomic prediction models and the entire pipeline for the Site-Frequency-Spectra analysis.

### Author Contributions

Funding acquired by GR, while the project was conceptualized by GR. Analyses conducted by NJ, JS, EN, SC, SW, BS, AO, DW and draft written by NJ and GR and reviewed by all authors.

### Large Language model statements

Language editing was supported using ChatGPT (OpenAI), a large language model (GPT-5.3-mini). The model was used for language clarity and grammar correction. No scientific content, analyses or interpretations was generated by the model. All scientific content were produced by authors.

## Supplementary

**Figure S1:**
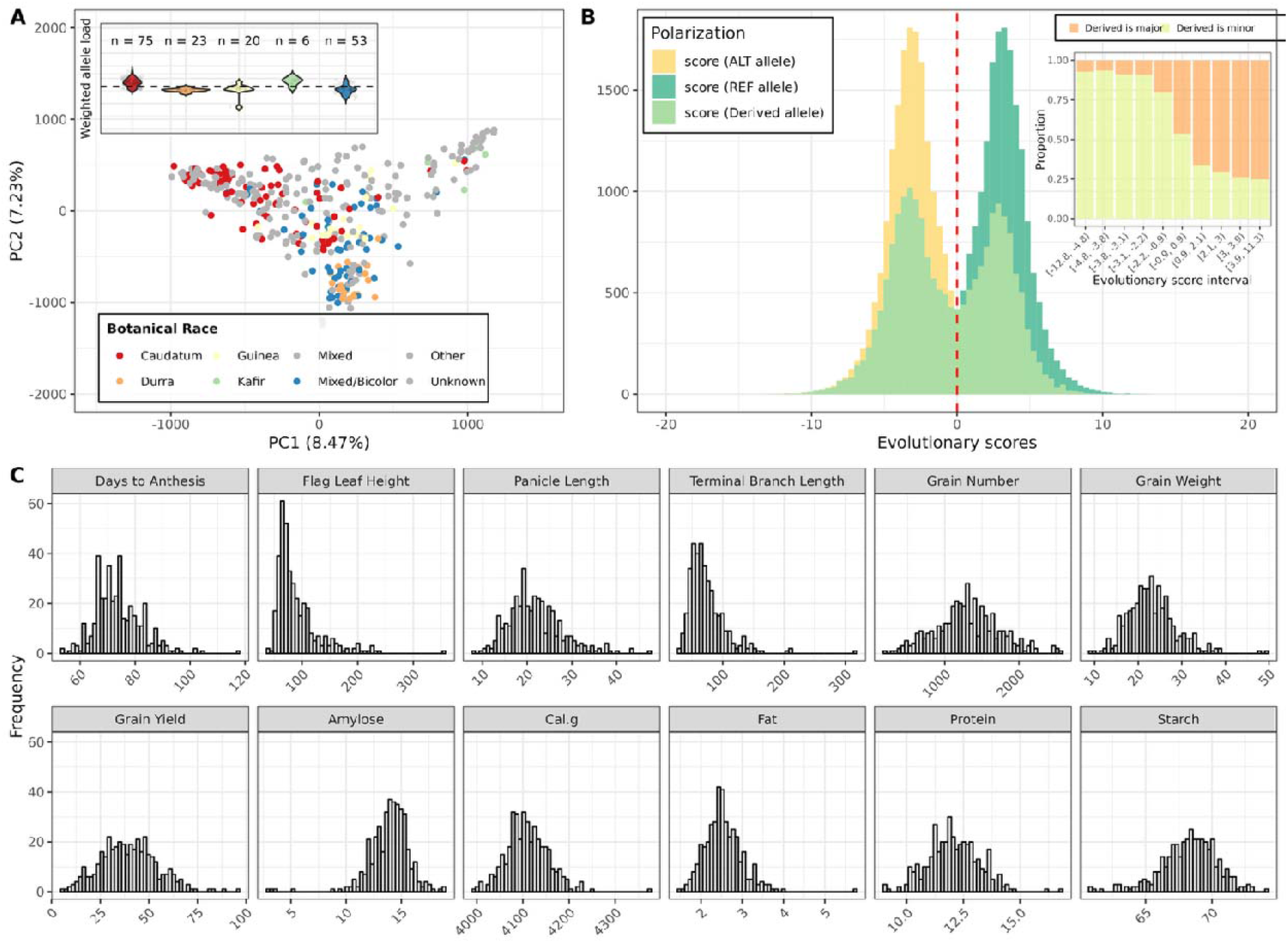
Phenotypic and genotypic data. **(A)** Principal component analysis (PCA) of the SAP diversity panel based on genome-wide SNP data. Points represent individual accessions, with colors indicating the botanical race. The violin plot shows the distribution of weighted allele load of nonneutral alleles for each of the botanical race. The stippled line indicates the average weighted load across all accessions, sample size for each botanical race shown in the violin plot. (**B**) Distribution of evolutionary scores for 0-fold degenerate sites with ancestral allele annotations. Evolutionary scores, as estimated with the Protein-Language-Model (PLM) ESM2, shown for three different polarization methods (i) Scores shown so the reflect the probability of observing the derived allele (DA) relative to the ancestral allele (AA), (ii) Scores polarized to reflect the probability of observing the reference allele (REF) relative to the alternative allele (ALT), and (iii) Scores polarized to reflect the probability of the ALT allele relative to the REF allele. The embedded plot shows, for each SNP-category, the proportion of derived alleles that is observed as major and minor allele. (**C**) Distribution of phenotypes for all traits included in the analysis.

**Figure S2:**
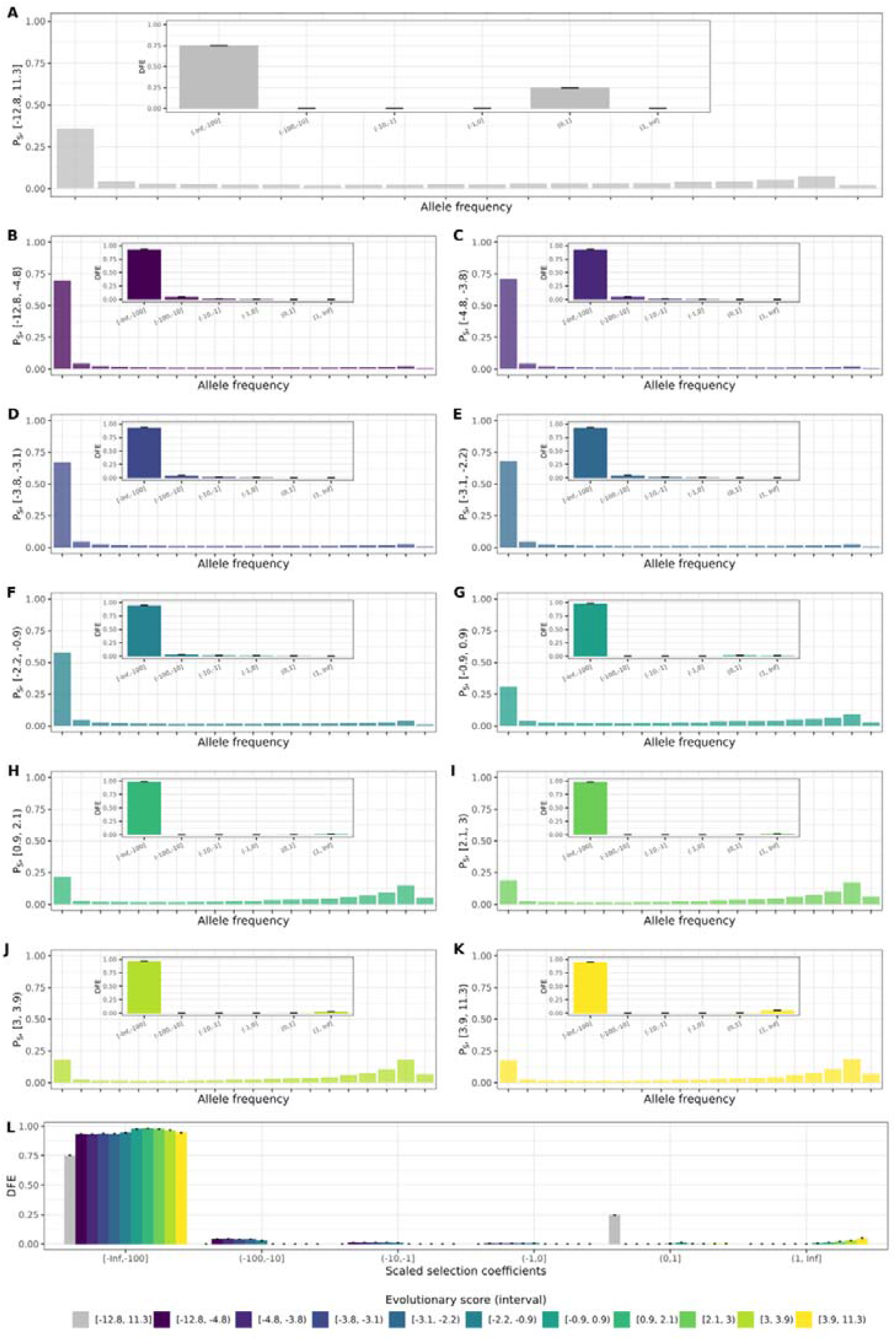
Predicted unfolded Site-Frequency-Spectrum (uSFS) shown for each of the ten SNP categories, in addition to their associated Distribution of Fitness effects (DFE). (**A**) The uSFS and corresponding DFE inferred from all 0-fold degenerate sites with ancestral allele annotations, the DFE plot is embedded. (**B-K**) The uSFS and underlying DFEs are shown for each of the ten mutation categories. mutation categories were formed by partitioning sites according to evolutionary score intervals estimated using ESM2. The lower and upper bounds of the evolutionary score intervals defining each category are indicated on the y-axis. (**L**) Summary of the DFEs across all ten mutation categories, shown together with the DFE inferred from the unpartitioned set of 0-fold degenerate sites.

**Figure S3:**
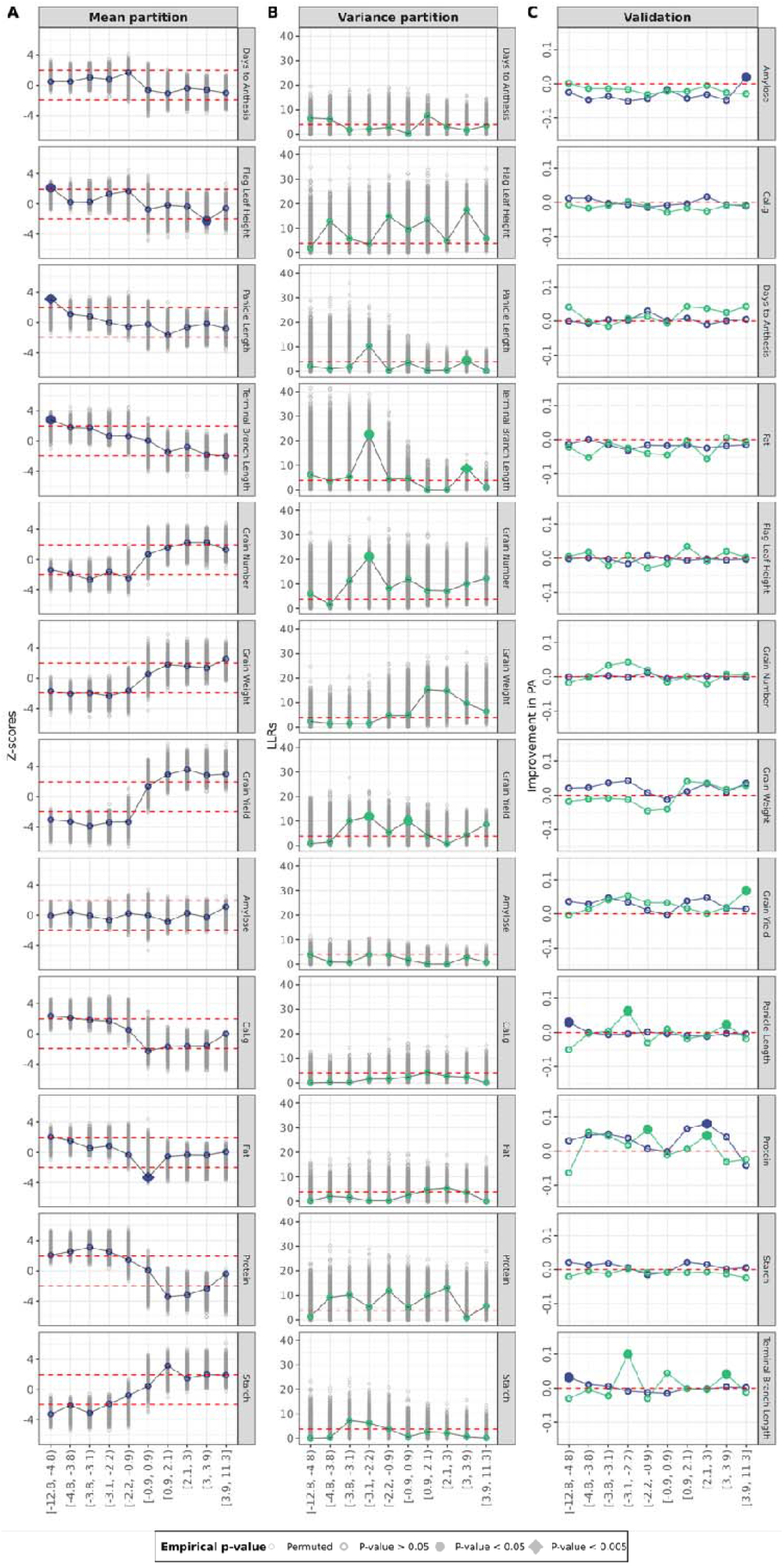
Impact of functional prioritization of variants, based on protein language model predictions, on genomic prediction model performance. (**A**) Improvement in prediction ability (PA) with the extended model M1 (mean partition) and M2 (Variance partition). Empirical significance derived from permutation tests are indicated by point shape. **(B)** Mean partition: **mutation** categories given as intervals of evolutionary scores for the derived allele shown on the x-axis. Points represent the estimated average effect of prioritized alleles standardized as Z-scores. Red dashed lines indicate the analytical significance threshold (p = 0.05). (**C**) Variance partition: Differences in log-likelihood ratios () between the baseline model M0 and extended model M2 are shown. Red dashed line indicates analytical significance (based on LLR-test). Empirical significance derived from permutation tests are indicated by point shape.

**Figure S4:**
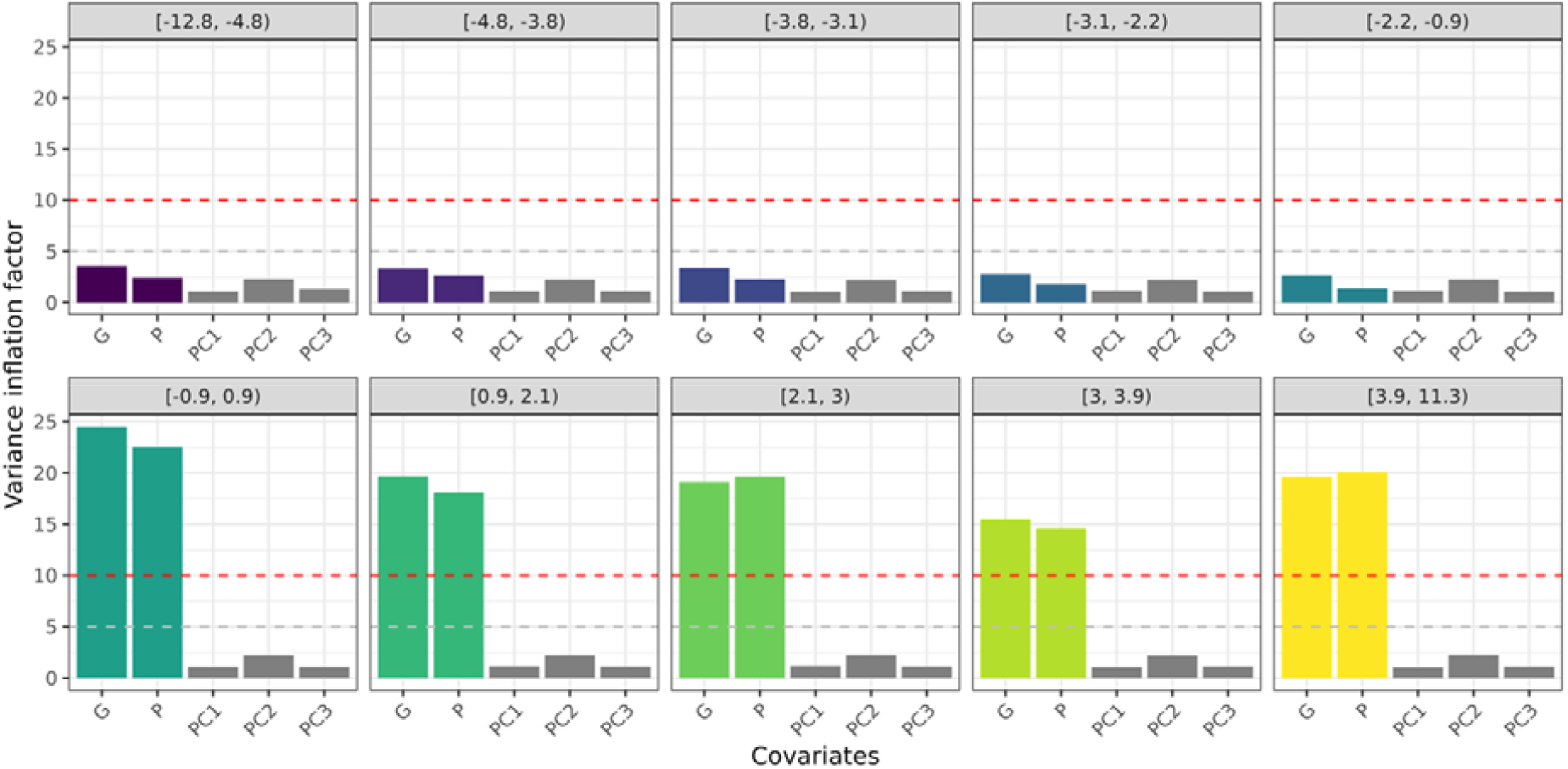
Variance Inflation factor for fixed effects in the mean partition models. Variance inflation factors (VIF) were calculated to assess multicollinearity among fixed effects in the mean partition model (M1). The genome-wide load (G) represents the total load of derived alleles across all 0-fold degenerate sites alignable to outgroups, while the P is the mutation load for a given mutation category, i.e., total count of derived alleles with evolutionary scores with a predefined interval. PC1-PC3 represent the first three principal components from a PCA analysis. Low VIF values indicates that the covariates show little collinearity, while high values indicate increased collinearity. Horizontal stippled lines indicate common threshold values for collinearity, with the dashed line at VIF = 5 and a line at VIF = 10. Values below 5 suggest low multicollinearity, while value between 5 and 10 indicate moderate collinearity and values above indicate high collinearity.

